# HPF1 REGULATES THE FORMATION OF FUS-DEPENDENT COMPARTMENTS BY PARP1 AND PARP2 ACTIVATION ON DAMAGED DNA

**DOI:** 10.1101/2025.09.01.673448

**Authors:** Anastasia S. Singatulina, Maria V. Sukhanova, Loic Hamon, Mikhail M. Kutuzov, David Pastre, Olga I. Lavrik

## Abstract

FUS participates in the formation of biomolecular condensates associated with PARP1-dependent synthesis of poly(ADP-ribose) (PAR). HPF1 regulates auto- and hetero-PARylation activities of PARP1 and PARP2 and may influence the formation of FUS compartments during PARP1 or PARP2 auto-PARylation. In this study, we used atomic force microscopy in combination with biochemical assay to investigate the formation of FUS compartments under activation of PARP1 and PARP2, when HPF1 modulates their activity. Similar to PARP1, FUS and PARylated PARP2 form DNA-rich compartments, indicating that PARP2 PARylation is sufficient for the formation of such compartments. The excess of HPF1 over PARP1 diminishes PARP1 activity and reduces the size of DNA-rich compartments. However, an excess of HPF1 over PARP2 does not significantly affect PARP2 activity and the size of compartments. Furthermore, HPF1 stimulates hetero-PARylation of FUS; this modification is stronger with PARP2 than with PARP1. HPF1-dependent intensive PARylation of FUS catalyzed by PARP1 or PARP2 impairs the assembly of DNA-rich compartment. These data provide a basis for investigating the effect of HPF1 on the formation of PAR-dependent condensates involving RNA-binding proteins like FUS, which interact effectively with PAR and show the ability to be targets of PARylation to regulate condensate formation at DNA damage sites.

**GRAPHICAL ABSTRACT:** 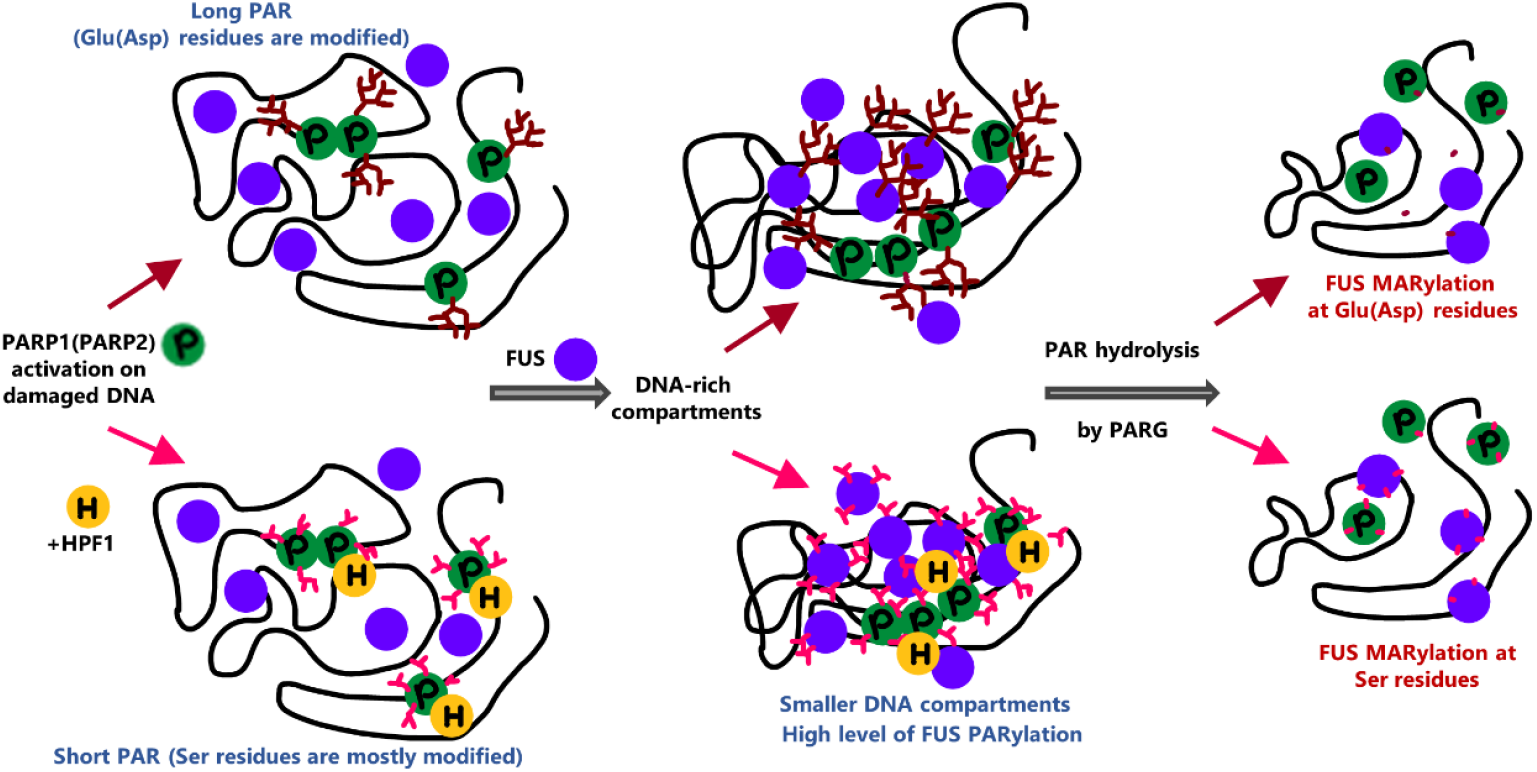

## INTRODUCTION

Poly(ADP-ribosyl)ation is one of the fastest cellular responses to DNA damage. Poly(ADP-ribose) polymerases 1 and 2 (PARP1 and PARP2), using NAD^+^ as a substrate, synthesize poly(ADP-ribose) (PAR) in response to DNA damage (1). This polymer is covalently modified of PARP1 and PARP2 themselves as well as a number of proteins under genotoxic stress(1). Degradation of PAR polymer is mediated by the enzymatic activity of poly(ADP-ribose) glycohydrolase (PARG)(2). An efficient and specific PARylation catalyzed with PARP1 seems to play an important role in attracting DNA repair proteins to DNA damage sites and the formation of PAR-dependent compartments for the following processing of DNA repair (1, 3, 4, 5). We have shown that autoPARylation of PARP1 in the presence of cellular cations is playing an important role in the initiation of compartment assembly on the DNA damage and stimulates binding DNA repair factors to PAR covalently linked to PARP1 (6, 7). This assumption was supported by other data published recently (8). PAR-hydrolyzing enzymes like PARG can dissociate compartments formed with PARylated PARP1, which could result in turnover of the DNA repair process (7, 9).

PARP2 is a homolog of PARP1 catalyzing DNA damage-dependent synthesis of PAR in the nucleus, but it is less active and contributes to only 15-25% of overall PAR synthesis in the cellular response to DNA damage (1, 10, 11). Similar to PARP1, PARP2 PARylation can also form a local signal at a DNA damage site and attract repair proteins through noncovalent interaction with PAR (10, 12). In PARP1^-/-^ cells, less active PAR synthesis catalyzed by PARP2 is sufficient for recruitment of DNA repair scaffold XRCC1 protein to chromatin after oxidative stress (12). However, the ability of PARP2 and PARP1 in the presence of histone PARylation factor 1 (HPF1), a factor regulating PARP1(2) catalytic activity, to perform the compartments has not been investigated. Indeed, HPF1 was recently identified as a key modulator of PARP1(2) activity by forming a transient joint active site with these proteins (13, 14, 15). The overall mechanism of the interaction between HPF1 and PARP1(2) is currently under intensive investigation (16, 17, 13, 18, 19, 20, 21) [;]. HPF1 has been demonstrated to change the PARP1(2)-mediated PARylation from glutamate/aspartate residues to serine residues, which regulates both auto-modification of PARP1 and PARP2 and hetero-modification of histones (17, 19, 20, 21). Furthermore, PARP1(2) interaction with HPF1 also leads to both stimulation and inhibition of PAR synthesis dependent on molar ratios of HPF1 to PARP1(PARP2) (18, 19, 20). Structural and biochemical investigation of PARP1(2) complex with HPF1 has shown that HPF1 blocks the histidine residues (His826 and His381) responsible for catalysis of PAR elongation in the active sites of PARP1 and PARP2, respectively, and thereby prevents PAR elongation, which leads to the formation of shorter polymers during protein PARylation (13, 18, 19). However, HPF1 stimulates the initiation of PARylation and does not inhibit elongation of PAR when the concentration of HPF1 is close to that of PARP1(2) or lower (18, 20). Studying the effect of HPF1 on the length and amount of PAR and protein hetero-PARylation is of great interest because these factors may influence chromatin relaxation, the recruitment of repair factors, and the interactions of the non-histone proteins with damaged DNA or PAR (1, 22, 23, 24, 25, 26, 27), thereby regulating further DNA repair and compartment formation.

Recently, a number of RNA-binding proteins prone to phase separation have been identified that colocalize at sites of DNA damage following PARP1 activation (28, 29, 30, 31). FUS/TLS (fused in sarcoma/translocated in liposarcoma) is precisely one of these proteins, and it has multiple cellular functions (32). FUS, together with EWS and TAF15, is a member of the FET family and one of the most abundant nuclear RNA-binding proteins that is highly PARylated under DNA damage (33, 34, 35). In addition, FUS interacts with PAR and undergoes phase separation (9, 29, 30, 31, 36). Recent studies suggest that FUS may be involved in DNA repair through the formation of biomolecular condensates, interacting with DNA damage signaling molecules such as PAR (4, 5, 9, 37, 38, 39, 40, 41). In addition, FUS could stimulate the repair process by inducing a temporary “compartmentalization” of damaged DNA (9). In this case, PARG-dependent PAR hydrolysis was demonstrated to disrupt DNA-rich compartments and likely promotes the release of FUS, DNA repair proteins, and repaired DNA (9). Such rapid dissociation of FUS-PAR compartments may be required to resume performing turnover of DNA repair (5, 9).

In this work, using an original approach based on a single-molecule analysis of PARylated PARP1 by AFM(9), we investigated the effect of HPF1 on PARP1(2) activity and the formation of compartments with participation of FUS. We performed a series of experiments using AFM to test the formation of FUS compartments involving PARylated PARP2 alone in the absence and the presence of HPF1. We investigated the impact of HPF1 on the PARylation of FUS by PARP1 and PARP2, considering their subsequent condensation with PARylated PARP1(2). We showed that FUS addition can support the assembly of the DNA-rich compartments in mixtures of damaged DNA and PARylated PARP2. We also observed that an increase in the concentration of HPF1 led to a decrease in the size of the emerging compartments formed with the participation of PARylated PARP1 and FUS, but at the same time, the number of the compartments remained at a similar level. PAR synthesis and compartment formation by PARP2 with HPF1 followed the same trend, but we observed a slight stimulation of PAR synthesis and compartment formation within an equimolar ratio of HPF1 to PARP2. However, the assembly of these compartments was impaired when FUS was added before the addition of NAD^+^ for PARP1(2) activation. A high level of FUS PARylation was observed in these conditions, particularly in the presence of HPF1. This FUS modification was observed to be stronger with PARP2 than with PARP1. These results demonstrate the effect of HPF1 on FUS PARylation. Hetero-PARylation of FUS is likely to hinder the formation of compartments containing FUS, PARylated PARP1 or PARP2, and damaged DNA. This provides a basis for further research into the impact of HPF1 on PAR-dependent condensate formation through hetero-PARylation of FUS that may be relevant in the context of searching for PARP inhibitors to treat diseases such as cancer, which is linked to so-called condensopathies (42, 43).

## MATERIALS AND METHODS

### Chemicals

Radioactive [α-^32^P]-ATP was produced in the Laboratory of Biotechnology at ICBFM (Siberian Branch of Russian Academy of Sciences [SB RAS], Novosibirsk, Russia). NAD^+^ and β-nicotinamide mononucleotide were purchased from Sigma-Aldrich (United States, catalog # 481911 and N3501, respectively), reagents for buffer and electrophoresis components from Sigma-Aldrich, United States (Tris, catalog # T6791; BSA, catalog # A9418; EDTA, catalog # E5134; HEPES, catalog # H3375), PanReacAppliChem, Germany (acrylamide/bis-acrylamide, catalog # A1089/A3636; urea, catalog # A1049), Molecular Group (DTT, catalog # 19733320), and Merk (NaCl, catalog # 106404). Olaparib (AZD2281, Ku-0059436) was purchased from Apexbio Technology (Cat#A4154k).

### Plasmids

#### Protein Expression and Purification

Recombinant human PARP1 and PARP2 (Table 1) were expressed in insect *Trichoplusia ni High Five*™ (Hi5) cells and purified as previously described (45). Recombinant bovine PARG (Table 1) was expressed in BL21(DE3) *Escherichia coli (E. coli)* cells and purified as previously described (46). Recombinant human HPF1 (Table 1) was expressed in Rosetta (DE3) *E. coli* cells and purified as previously described (20). Full-length His-tagged FUS(1-526), His-tagged FUS(1-526), His-tagged FUSΔLCD(164-526), His-tagged FUSΔRGG1(1-455), His-tagged FUSΔRGG1-2(1-375), His-MPB-tagged FUS-6E (S26E, S30E, T68E, S84E, S87E, S117E), and His-MPB-tagged FUS-12E (T7E, T11E, T19E, S26E, S30E, S42E, S61E, T68E, S84E, S87E, S117E, S131E) (Table 1) were expressed in BL21(DE3) *E. coli* and purified as previously described (9). APE1 (Table 1) (apurinic/apyrimidinic endonuclease 1) was expressed in BL21(DE3) *E. coli* and purified as previously described (47).

**Table 1.** Recombinant DNA was used in this study.

| Recombinant DNA | Source | Identifier |
| --- | --- | --- |
| Vector pBR322 | New England BioLabs | Cat#N3033S |
| pET-FUS | Singatulina et al., 2019 (9) | N/A |
| pET-FUS-ΔLCD | Singatulina et al., 2019 (9) | N/A |
| pET-FUS_1-374 | Singatulina et al., 2019 (9) | N/A |
| pET-FUS_1-454 | Singatulina et al., 2019 (9) | N/A |
| pET-MBP-FUS_FL_6E | Monahan et al., 2017 (44) | Addgene #98652 |
| pET-MBP-FUS_FL_12E | Monahan et al., 2017 (44) | Addgene #98655 |
| pLate31-HPF1-His | Kurgina et al., 2021 (20) | N/A |
| pGEX-2T-bPARG-GST | Dr. V. Schreiber | N/A |
| pXC53-hAPE1 | Dr. S.H. Wilson | N/A |
| pVL-mPARP-2-F | Dr. J.-C. Amé | N/A |
| pVL 1392-hPARP1 | Dr. J.-C. Amé | N/A |

Yeast nicotinamide mononucleotide adenylyltransferase (NMNAT) was kindly provided by Dr. S.I. Shram (Institute of Molecular Genetics, Russian Academy of Science, Moscow, Russia).Bovine poly(ADP-ribose) glycohydrolase (PARG) was kindly provided by Dr. T.A. Kurgina (ICBFM, Novosibirsk, Russia). Human APE1 was kindly provided by Dr. N.S. Dyrkheeva (ICBFM, Novosibirsk, Russia).

SDS-PAGE was used to monitor the purity of the proteins and the proteolysis step at all stages of purification.

### DNA Substrates

The pBR322 plasmid containing DNA breaks (damaged plasmid DNA) was prepared using heat and acid treatment to create abasic sites followed by apurinic/apyrimidinic site cleavage with APE1 activity as previously described (9).

### Preparation of samples for atomic force microscopy

Protein complexes for AFM analysis were formed in reaction mixtures (20 μL) containing binding buffer (12.5 mM HEPES pH 7.6, 12.5 mM NaCl, 5 mM MgCl_2_, 1 mM DTT, 100 mM urea), 30 nM PARP1(2), 400 nM FUS, 12.5 nM damaged DNA (damaged pBR plasmid), and 30-960 nM HPF1 as indicated in the figure legends.

To image PARylated proteins or compartments, 30 nM PARP1(2) was incubated with 12.5 nM damaged pBR, 0.3 mM NAD^+^ in the absence or presence of FUS (400 nM), and 30-960 nM HPF1 in the binding buffer at 37°C for 5 min. The reaction mixtures were incubated with 400 nM FUS as indicated at 37°C for 1 min, then diluted 10-fold in the binding buffer containing 200 mM urea and immediately deposited on mica.

To analyze the dissociation of compartments in the presence of PARG, the reaction mixtures were pre-incubated with 400 nM FUS at 37°C for 1 min in the reaction buffer, followed by incubation with 40 nM PARG at 37°C for 30 min. After that, the reaction mixtures were diluted 10-fold in the binding buffer containing 200 mM urea and immediately deposited on mica.

To adsorb the molecules on mica, putrescine was added to the solution to a final concentration of 1 mM, after which a 10 μL droplet was deposited on the surface of freshly cleaved mica at room temperature for 30 s and dried for AFM imaging as described previously (9).

### AFM imaging and image analysis

AFM images were recorded in air by using a Nanoscope V Multimode 8 (Bruker, Santa Barbara, CA) in PeakForce Tapping (PFT) mode using Scanasyst-Air probes (Bruker). Continuous force-distance curves were thus recorded with an amplitude of 100-300 nm at low frequency (1-2 kHz). PFT mode decreases the lateral and shear forces. Images were recorded at 2048 × 2048 pixels at a line rate of 1.5 Hz.

The “Section” tool in the Nanoscope Analysis software (version 1.70) was used to determine the molecular dimensions of the protein particles and compartments. Cross-sections of protein particles were made, and the diameter (d) of each single particle was measured.

### Radioactive assay of protein PARylation and PAR hydrolysis by PARG in vitro

[^32^P]-NAD^+^ labeled on the adenylate phosphate was synthesized in a reaction mixture (100 μL) containing 2 mM β-nicotinamide mononucleotide, 1 mM ATP, 0.25 mCi of [α-^32^P]-ATP (1000 Ci/mmol), 1.5 mg/mL nicotinamide mononucleotide adenylyl transferase (NMNAT), 25 mM Tris-HCl (pH 7.5), and 20 mM MgCl_2_ was incubated for 1 h at 37°C. The enzyme was denatured at 65°C for 10 min, and precipitated proteins were removed by centrifugation.

An in vitro poly(ADP-ribosyl)ation assay was performed in the reaction mixtures (20 μL) containing binding buffer (12.5 mM HEPES pH 7.6, 12.5 mM NaCl, 1 mM DTT, 100 mM urea, 5 mM MgCl_2_), 12.5 nM damaged DNA, 30 or 250 nM PARP1(2), 30-1000 nM HPF1, 0.3 mM NAD^+^, 0.4 μCi [^32^P]NAD^+^ and 1 µM FUS or FUS mutants as indicated in the figure legends. The reactions were initiated by the addition of NAD^+^. The reaction mixtures were incubated at 37°C for 15 min for PARP1 and 30 min for PARP2 at 37°C and stopped by adding SDS-sample buffer and heating for 5 min at 90°C. The reaction mixtures were analyzed by 10% SDS-PAGE with subsequent phosphorimaging and/or colloidal Coomassie staining.

For the analysis of the PARylation of PARP1(2) after PARG treatment, 30 nM PARP1(2), 400 nM FUS, 12.5 nM damaged pBR plasmid, and 30-960 nM HPF1 (as indicated in the figure legend) were incubated in the binding buffer followed by reaction initiation by the addition of 0.3 mM NAD^+^. The reaction mixtures were incubated at 37°C for 15 min for PARP1 and 30 min for PARP2 at 37°C and then were stopped by Olaparib to a final concentration of 250 nM. Then, PARG was added to a final concentration of 40 nM, after which the mixture was incubated for 3-24 minutes at 37°C. The reaction mixture (20 μL) was stopped by adding 4 μL of the loading solution containing 90% formamide, 50 mM EDTA, 0.1% xylene cyanol, and 0.1% bromophenol blue, heated for 5 min at 95°C, and the products were separated by denaturing electrophoresis in 10% polyacrylamide gel followed by visualization with phosphorimaging.

### Quantification and Statistical Analysis

AFM analysis: images shown in the figures are representative of three different and independent samples. The “particle analysis” tool in the Nanoscope Analysis software (version 1.50) was used to determine the diameter of the adsorbed molecules and multicomponent complex particles from at least three independent samples. For each particle of interest, the particle analysis tool measured the maximum diameter of the particle. Significance of height was tested by using a t-test: *, p < 0.05; **, p < 0.01; ***, p < 0.001; ****, p < 0.0001; ns, non-significant. The particle diameter is twice the smallest radius of a circle in which the particle can be placed.

PAGE analysis: bands of [^32^P]-PAR-labeled PARP1, PARP2, FUS, and its mutants were quantified by means of the Quantity One Basic software (Bio-Rad).

## RESULTS

### FUS forms DNA-rich compartments when PARP1 is activated in the presence of HPF1

We previously showed that FUS and PARP1 PARylated on damaged DNA form complex supramolecular structures (compartments) in which damaged DNA is concentrated while excluding undamaged DNA(9). The interaction of FUS with PAR plays an important role in the formation of compartments (30, 9, 31, 36, 48, 49, 50); furthermore, the structure of PAR and the level of its synthesis can significantly influence this process (36, 50). It was shown that HPF1 affects the autoPARylation of PARP1 and PARP2 in the presence of activated DNA, short DNA duplexes, or nucleosomes (16, 18, 19, 20, 21). The stimulation of PARP1 and PARP2 activity was detected in a defined range of HPF1 and NAD^+^ concentrations at which there is no HPF1-dependent increase in the hydrolytic NAD^+^ consumption (19, 20). PARP2 was found to be stimulated more efficiently by HPF1 than PARP1 in autoPARylation and histone hetero-PARylation (20, 21). Therefore, we hypothesized that the presence of HPF1 will affect the assembly of the FUS compartments formed during PARP1 PARylation. First, we analyzed PARP1 activation in the presence of HPF1 using a circular damaged pBR322 plasmid (hereafter referred to as damaged DNA) as a substrate and measured the size of PARylated molecules using AFM imaging (Figure 1A (upper panels)). We observed that the size of PARylated PARP1 molecules decreased when the concentration of HPF1 increased (Figure 1B). That is in agreement with the previous data showing that an excess of HPF1 over PARP1 stimulates initiation of PAR synthesis but blocks its elongation (18, 19, 20). For HPF1-independent PAR synthesis, the PARylated PARP1 molecules had an average size of around 34 nm. In the presence of HPF1, the molecular size was reduced to 23– 29 nm (Figure 1B). Thus, the size of the synthesized PAR in the case of PARP1 autoPARylation decreased with the addition of HPF1, ranging from an equimolar ratio of HPF1:PARP1 to a higher ratio of HPF1:PARP1 (16:1). The further measurements of size of PARylated molecules were not possible due to the high degree of crowding of the molecules on the AFM mica surface.

**Figure 1.**
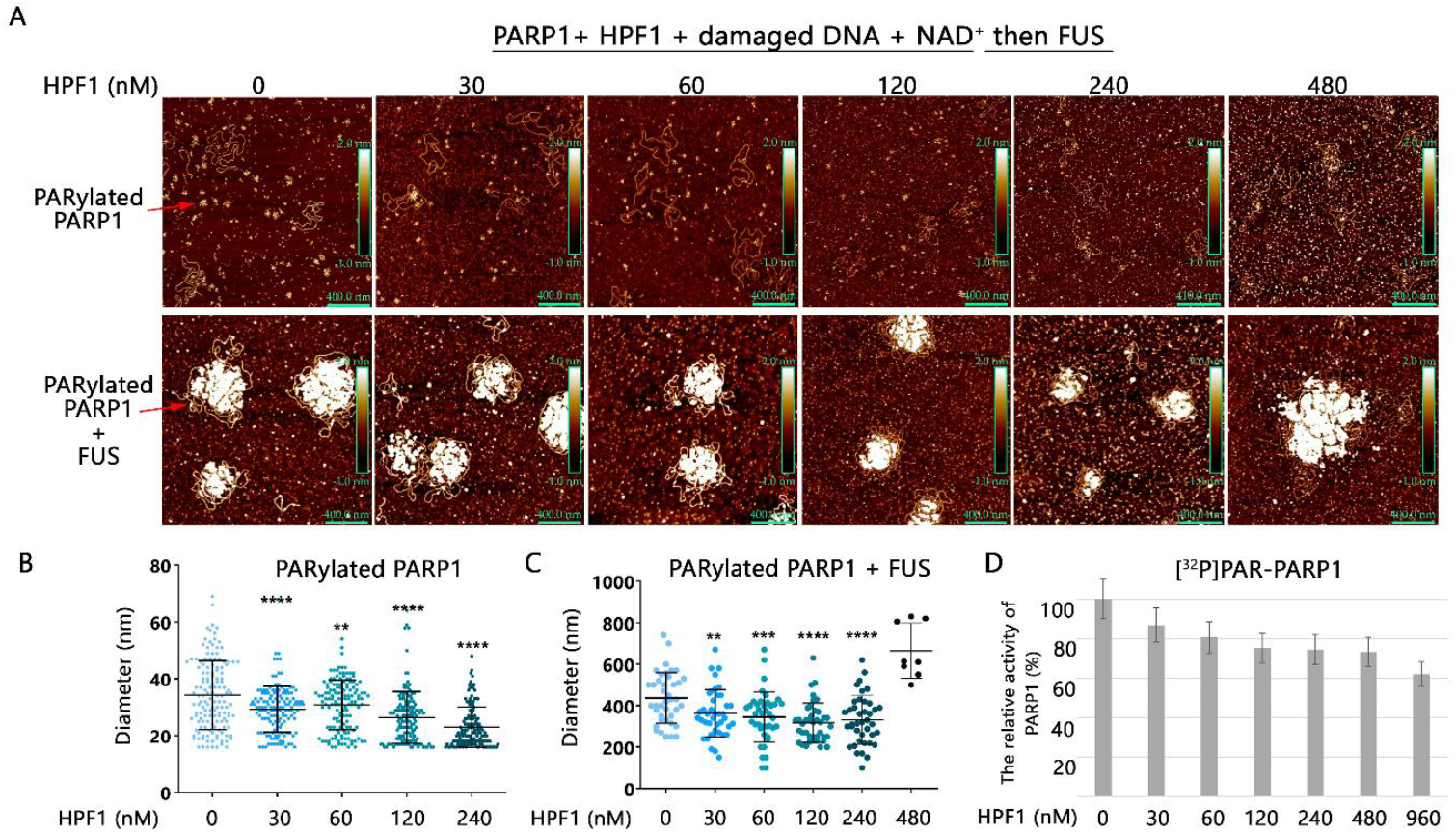
FUS forms DNA-rich compartments upon PARP1 activation in the presence of HPF1. (**A**) Upper panels: AFM images of PARylated PARP1 (30 nM) after incubation with damaged pBR plasmid (12.5 nM) and HPF1 (30-480 nM) for 5 min in the presence of NAD^+^ (300 μM). Lower panels: same conditions as on the upper panel, but followed by the addition of FUS (400 nM) and incubation for 1 min before deposition on a mica surface. Scale bar: 400 nm; Z scale: 2 nm. (**B**) Diameter of PARylated molecules were measured in images shown in (A, upper panels). **, p < 0.01; ***, p < 0.001; ****, p < 0.0001, paired t-test; n = 140 per sample. Horizontal bars indicate mean; scanned area: 20 μm^2^ per sample, 3 samples per condition. (**C**) Diameter of compartments were measured in images shown in (A, lower panels). **p < 0.05, paired t-test; n = 40 per sample (except 480 nM HPF1). Horizontal bars indicate mean; scanned area: 30 μm^2^ per sample, 3 samples per condition. (**D**) Diagrams showing calculation of the relative level of PARP1 activity in the presence of HPF1 after separation of the products by 10% SDS-PAG? (Figure S2A). PARP1 (30 nM) was incubated with damaged DNA (12.5 nM), 0.3 mM NAD^+^, and 0.4 μCi [^32^P]-NAD^+^ in the absence or presence of HPF1 (30-960 nM). The relative level of PAR synthesis was normalized to the level of PAR synthesis catalyzed by PARP1 alone for 15 min without HPF1.

Further, we investigated the role of HPF1 in the formation of damaged DNA-containing compartments formed by the interaction of FUS with PARylated PARP1. Previous studies using AFM showed local recruitment of FUS to PAR synthesized by PARP1 at damaged DNA sites and the formation of large compartments in which damaged DNA is enriched (9). Here, we used the same conditions to observe compartment formation, but the activity of PARP1 was modulated by HPF1. For compartment formation, FUS was added after PARP1-dependent synthesis of PAR in the absence or presence of HPF1, and compartments were formed over the entire range of HPF1 concentration (30-480 nM) (Figure 1A (lower panels)). The size of the compartments decreased, as well as the size of the PARylated PARP1 molecules (Figure 1B), except for the compartments formed in the presence of 480 nM HPF1 (Figure 1C), where only a few very large compartments were formed, most likely having a denser structure (Figure 1A (lower panels)). AFM imaging of pBR, PARP1, and FUS in the presence or absence of HPF1 showed that compartments were not formed in the absence of NAD^+^ (Figure S1).

Next, we measured PARP1 activity using an assay of PAR synthesis to estimate the amount of [^32^P]-labeled ADP-ribose polymer synthesized by using [^32^P]-NAD^+^ in the presence of different concentrations of HPF1, which we used for AFM analysis (Figure 1D and Figure S2A). An excess of HPF1 (60-960 nM) over PARP1 (30 nM) caused a decrease of PAR synthesis (Figure 1D).

In general, we observe a decrease in the amount of synthesized PAR, the size of PARylated PARP1 molecules, and, as a consequence, the size of FUS-DNA-rich compartments formed in the presence of HPF1 (Figure 1).

### PARP2 activation enables the formation of DNA-rich compartments with theparticipation of FUS, and HPF1 does not hinder this process

PARP2 is the closest homologue of PARP1 (1, 11). Like PARP1, PARP2 catalyzes the synthesis of poly(ADP-ribose) and plays an important role in DNA damage response (10, 12, 51, 52). PARP2 and PARP1 can perform similar functions within the cell but also demonstrate different biological activity (11). It is worth noting that in vitro studies have shown that PARP2 exhibits lower catalytic activity than PARP1 and that these two proteins have different affinities for specific types of DNA damage (53, 54, 55, 56). Although the functional relationship between FUS and PARP1 has previously been studied *in vitro* (9, 49, 50), the interaction between FUS and PARP2 has not. The interaction of FUS with PARylated PARP1 induces the formation of damaged DNA-rich compartments(9). In principle, the same effect can also be generated by PARylated PARP2. Similar to the experiments with PARP1 (Figure 1), we analyzed PARP2 activation in the absence or presence of HPF1 using damaged pBR plasmid as substrate and measured the size of PARylated molecules using AFM imaging (Figure 2A).

**Figure 2.**
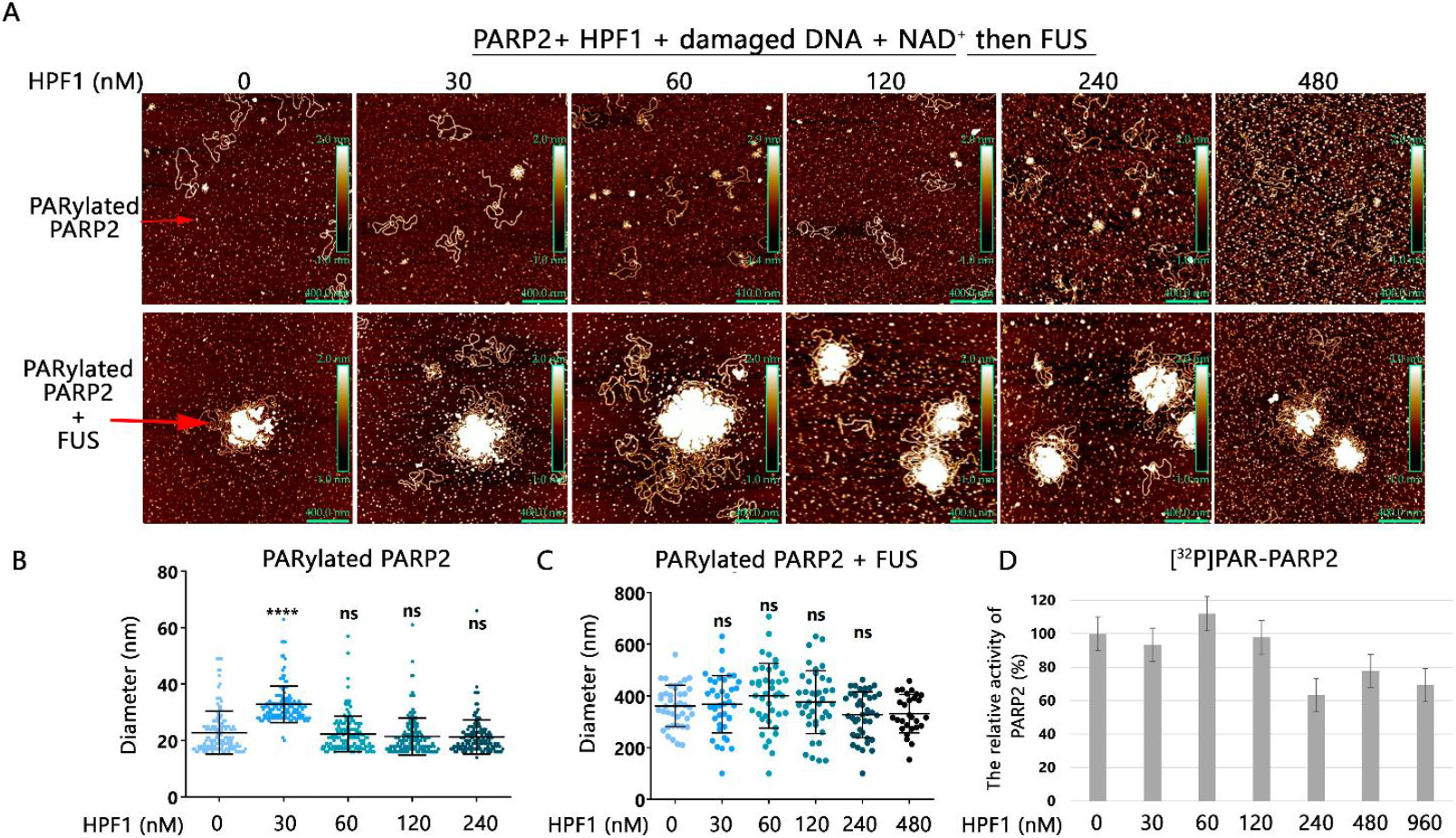
FUS forms DNA-rich compartments upon PARP2 activation in the presence of HPF1. (**A**) Upper panels: AFM images of PARylated PARP2 (30 nM) after incubation with damaged pBR plasmid (12.5 nM) and HPF1 (30-480 nM) for 5 min in the presence of NAD+ (300 μM). Lower panels: Same conditions as on the upper panel, but followed by the addition of FUS (400 nM) and incubation for 1 min before deposition on mica surface. Scale bar: 400 nm; Z scale: 2 nm. (**B**) Diameter of PARylated molecules was measured in images shown in A (upper panels). ****p < 0.0001, ns, non-significant, paired t-test; n = 140 per sample. Horizontal bars indicate mean; scanned area: 20 μm^2^ per sample, 3 samples per condition. (**C**) Diameter of compartments were measured in images shown in (A, lower panels). ns p > 0.05, paired t-test; n = 40 per sample (except 480 nM HPF1). Horizontal bars indicate mean; scanned area: 30 μm^2^ per sample, 3 samples per condition. (**D**) Diagrams showing calculation of the relative level of PARP2 activity in the presence of HPF1 after separation of the products by 10% SDS-PAG? (Figure S2B). PARP2 (30 nM) was incubated with damaged DNA (12.5 nM), 0.3 mM NAD^+^, and 0.4 μCi [^32^P]-NAD^+^ in the absence or presence of HPF1 (30-960 nM). The relative level of PAR synthesis was normalized to the level of PAR synthesis catalyzed by PARP2 alone for 30 min without HPF1.

For HPF1-independent PAR synthesis, the PARylated PARP2 had an average size of around 22 nm (Figure 2B). In the presence of HPF1, the size of the modified PARP2 increased to 36 nm when the ratio of HPF1 to PARP2 was equimolar (1:1) (Figures 2A (upper panels) and 2B). This fact is in agreement with the previously published data (20, 21). Unlike PARP1, the size of PARylated PARP2 molecules was slightly affected by an increase in HPF1 concentration (60-240 nM) that was across a range of molar ratios from 1:1 to 4:1 (Figures 2B).

Using AFM, we have also detected that FUS can form DNA-rich compartments after PARylation of PARP2 both in the absence and in the presence of HPF1 (Figure 2A (lower panel)). Nevertheless, the size of the compartments did not change significantly in the presence of HPF1, and the number of the compartments remained at the same level (Figure 2C). AFM measurements also showed that the size of PARylated molecules and FUS-DNA-rich compartments formed after PARP2 modification is only slightly affected at the HPF1:PARP2 molar ratio greater than 1:1 (Figure 2B, C). We also estimated PARP2 activity using a radioactive assay to measure the amount of ADP-ribose polymer synthesized under variation of HPF1 concentration (30-960 nM), which we used in the AFM assay (Figure 2D and Figure S2B). Excess of HPF1 (30-120 nM) over PARP2 (30 nM) in the range of 1:1 to 4:1 had a low effect on PARP2 autoPARylation (Figure 2D).

These data indicate that interaction between FUS and PAR-PARP2 can lead to the formation of DNA-rich compartments, and this process is slightly influenced by HPF1 when its molar concentration is equal to or in excess of PARP2 (Figure 2).

### PARG can disrupt the FUS compartments that form when PAR synthesis occurs in the presence of HPF1

It is known that PARylation is a reversible process of protein modification, primarily due to the activity of PARG, which catalyzes the cleavage of both free poly(ADP-ribose) and that attached to proteins (2). We have previously demonstrated that PARG-dependent degradation of PAR bound to PARP1 can disrupt FUS-PARylated PARP1 compartments (9). In vitro experiments have demonstrated that shorter PAR is degraded more slowly by PARG than longer PAR (57). Since an excess of HPF1 over PARP1 could decrease the length of PAR (Figure 1), we investigated the effect of PARG enzymatic activity on FUS compartments formed during PARP1 activation in the presence of HPF1 (Figure 3). AFM imaging revealed that PARG almost completely degrades the compartments formed by PARP1 after PAR synthesis in the presence of HPF1 and following the addition of FUS (Figures 3A and 3B). This suggests that PARG hydrolyzes the ADP-ribose polymer attached to PARP1 because PAR is the main inducer of DNA-rich compartment formation with the participation of FUS (9). Next, to detect PAR degradation under compartment formation conditions (Figures 1 and 2), we also used a polyacrylamide gel electrophoresis (PAGE)-based assay. This assay enables the detection of PAR (Figure S3).

**Figure 3.**
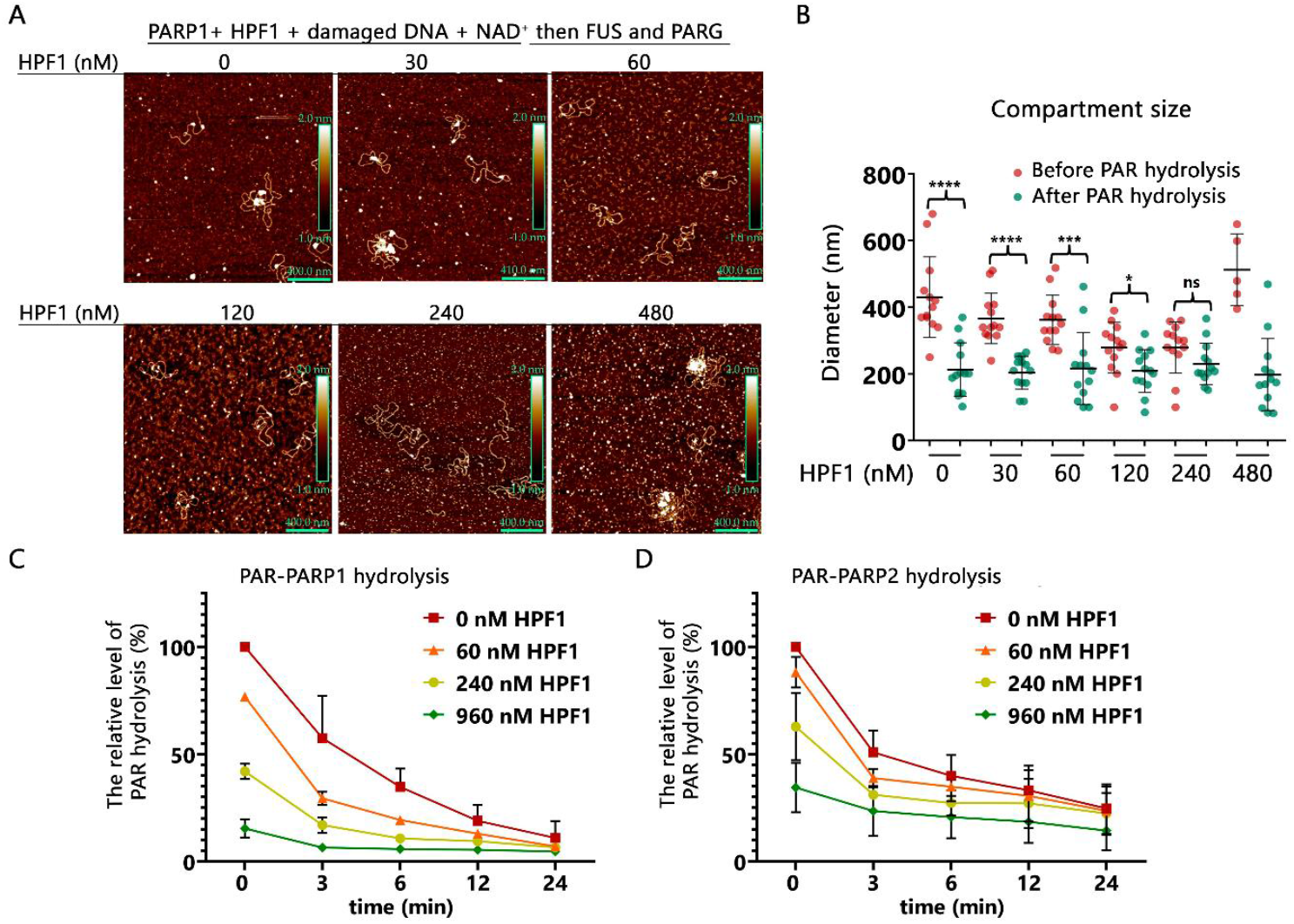
PARG was able to efficiently hydrolyze PAR synthesized by PARP1(2) in the presence of HPF1 under conditions of FUS compartment formation. (**A**) Representative AFM images of DNA-rich compartments formed by FUS after PARP1 activation in the absence or presence of HPF1 (30-480 nM) and subsequent hydrolysis of PAR by 40 nM PARG for 30 min. Scale bar: 400 nm; Z scale: 2 nm. AFM images of PARylated PARP1 (30 nM) after incubation with damaged pBR plasmid (12.5 nM) and HPF1 (30-480 nM) for 5 min in the presence of NAD^+^ (0.3 mM) and followed by the addition of FUS (400 nM) and incubation for 1 min before addition of PARG (40 nM). (**B**) The diameter of the compartments was measured in images obtained in conditions similar to Figure 1A (red circle, before PARG adding) and in images shown in A (green circle, 30 minutes after adding PARG). *, p < 0.05; ***, p < 0.001; ****, p < 0.0001; ns, non-significant, paired t-test; n = 13 per sample (at 0, 30, 60, 120, and 240 nM HPF1 in the absence or presence of PARG) and n = 6 per sample (480 nM HPF1 in the absence of PARG). Horizontal bars indicate mean; scanned area: 30 μm^2^ per sample, 3 samples per condition. (**?**) and (**D**) The graphs present quantification of [^32^P]-labeled PAR after PARG treatment (Figure S3). The relative level of PAR hydrolysis in the presence of FUS was normalized to the amount of PAR synthesized by PARP1 (**C**) or PARP2 (**D**) in the absence of HPF1 for 30 min.

Before the addition of PARG, analysis of ADP-ribose polymers produced by PARP1 and PARP2 revealed a reduction in polymer amount and length dependent on HPF1 concentrations (Figure 3C, D and Figure S3). PARP1 exhibited a more pronounced decrease in PAR amount and length than PARP2. This is consistent with the analysis of the protein PARylation level by SDS-PAGE in the presence of HPF1, which showed a decrease of up to 40% for PARP1 modification and up to 30% for PARP2 modification at the highest excess of HPF1 (32:1) over PARP1(2) (Figures 1D and 2D and Figure S2). Although FUS has been shown to partially protect PAR from hydrolysis under conditions of FUS compartment formation (9), PARG can efficiently hydrolyze PAR synthesized by PARP1(2) in the presence of HPF1 (30-960 nM) when FUS is added at a concentration that leads to the formation of compartments (Figure 3C, D).

Thus, PARG-dependent degradation of PAR facilitates the dissociation of damaged DNA-rich compartments, as evidenced by the appearance of isolated DNA molecules and small aggregates on the mica surface (Figure 3A). Although PARG cannot remove the first ADP-ribose units attached to a protein (58), its activity is sufficient to dissociate DNA-rich compartments in vitro, also in the case when PARP1(2) is PARylated in the presence of HPF1. Therefore, mono(ADP-ribosyl)ation of PARP1(2) does not contribute to the compartment formation (Figure 3A, B).

### Hetero-PARylation of FUS influences the formation of DNA-rich compartments

It has been observed that FUS is a substrate for PARylation by PARP1 both in the cells and in vitro (9, 34, 35, 49, 59). Furthermore, FUS effectively binds to PAR via its RGG domains, and these interactions play a significant role in FUS condensation (9, 30, 48, 36). Therefore, the PARylation of FUS by PARPs (the attachment of negatively charged ADP-ribose) could affect its binding to PAR and consequently influence the compartment formation by interaction of FUS with PAR. This suggests that the formation of FUS compartments has to be dependent on the level of FUS PARylation. We tested compartment formation when FUS and PARP1 were added simultaneously, followed by the addition of NAD^+^ to initiate PAR synthesis (Figure 4). Under these conditions, FUS was previously shown to be effectively PARylated (9, 49). Using AFM imaging, we observed a general decrease in compartment size when FUS was added before the addition of NAD^+^ for PARP1 activation. In this case, smaller compartments were formed (Figure 4B). Under such conditions, the part of the FUS protein appears to undergo PARylation by PARP1 and does not bind to PAR attached covalently to PARP1; therefore, we observed that the size of the compartments was decreased (Figure 4B). It was suggested that the level of auto-PARylation of PARP1(2) and hetero-PARylation of the target proteins can be influenced by HPF1, as it was previously demonstrated for core histones in nucleosome core particles (20, 21). Thus, HPF1 may affect PARP1(2)-mediated PARylation of FUS in vitro, thereby influencing compartment formation. To test this hypothesis, we examined compartment formation with AFM when FUS and HPF1 were added together before the addition of NAD^+^ for PARP1 activation (Figure 4A, B). In this case, the presence of HPF1 had no significant effect on the formation of FUS compartments or their size (Figure 4B).

**Figure 4.**
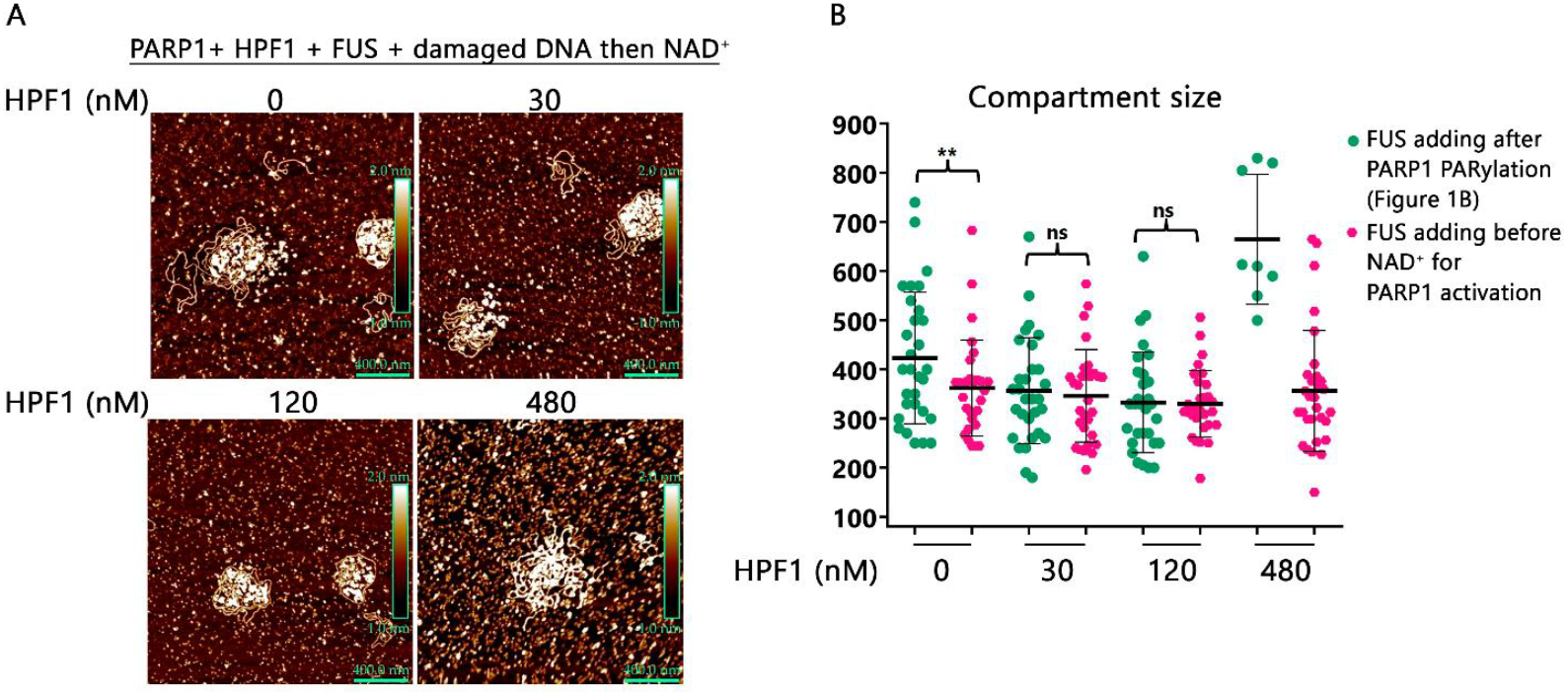
The size of DNA-rich compartments decreased if FUS was added before initiation of PARylation. (**A**) Representative AFM images of compartments formed by FUS during PARP1 activation under indicated conditions (same as in Figure 1A). FUS was added prior to NAD^+^, which was followed by the activation of PARP1. Scale bar: 400 nm; Z scale: 2 nm. (**B**) Comparing the size of compartments measured in images shown in Figure 1B (green circle) and images shown in A (pink circle); **p > 0.01; ns, non-significant; paired t test; n = 30 per sample (except 480 nM HPF1). Horizontal bars indicate mean; scanned area: 30 μm^2^ per sample, 3 samples per condition.

To explain how compartment size is regulated based on the order in which FUS is added, we hypothesized that low-level FUS PARylation occurs when FUS is added after PAR synthesis begins, while high-level FUS PARylation occurs when FUS is added before the addition of NAD^+^ for PARP1 activation. To challenge this hypothesis, FUS PARylation by PARP1 was examined in the presence or absence of HPF1 (Figure S4A). When FUS was added before the addition of NAD^+^ for PARP1 activation, a high level of FUS PARylation was observed. (Figure S4A). Adding HPF1 stimulates PARP1 activity, thereby increasing the level of FUS modification by ∼2.0-fold (Figures S4). Thus, FUS modification was strongly influenced by HPF1, as it increased the level of FUS hetero-PARylation catalyzed by PARP1 (Figure S4A). Therefore, HPF1 may play an important role in FUS-dependent compartment formation during PARP1 activation because it stimulates FUS hetero-PARylation (Figure S4) with the following disruption of its interaction with PAR.

FUS phase separation is determined by its ability to self-assemble and/or interact with PAR (9, 30, 31, 36, 48, 44). The N-terminal low complexity domain (LCD) and the C-terminal RGG(2,3) domains have been identified as critical for regulating FUS phase separation in PAR-containing systems (9, 30, 31, 44); LCD is not involved in PAR binding but mediates FUS self-assembly (31, 44). The RGG(2,3) domains in turn possess PAR binding activity (9, 30, 56, 48). Using quantitative proteomics, ADP-ribosylation at serine residues in the RGG1 and RGG2 domains of FUS was observed in HeLa cells in response to oxidative stress induced by hydrogen peroxide (60, 61). This suggests that FUS PARylation can potentially affect its phase separation, especially when its PARylation by PARP1(2) occurs in the presence of HPF1. We then set out to determine whether mutations in the different FUS domains affect its PARylation by PARP1 in the absence or presence of HPF1 (Figure 5). For these experiments, we used FUS phosphomimetics (FUS6E and FUS12E), containing substitutions of 6 or 12 serine/threonine residues with glutamine in the FUS LCD domain, or FUS mutants with deletion of the N-terminal LCD (FUSΔLCD) or C-terminal RGG2,3 domains (FUSΔRGG3 and FUSΔRGG2,3) (Figure 5A). FUS LCD phosphorylation was shown to impair its phase separation, thereby suppressing the formation of damaged DNA-rich compartments (9, 44, 62). We observed that HPF1 stimulated PARylation of all FUS mutants by PARP1 (Figure 5B, C). There was no significant difference in the protein modification in the comparison between the FUSΔLCD and FUS wild-type. By contrast, mutations in the LCD domain or the deletion of one or two RGG domains reduced FUS PARylation in both the absence and presence of HPF1 (Figure 5C). The data suggest that deletion of at least two RGG domains in FUS significantly reduces its PARylation by PARP1, whether HPF1 is present or not (Figure 5B, C)

**Figure 5.**
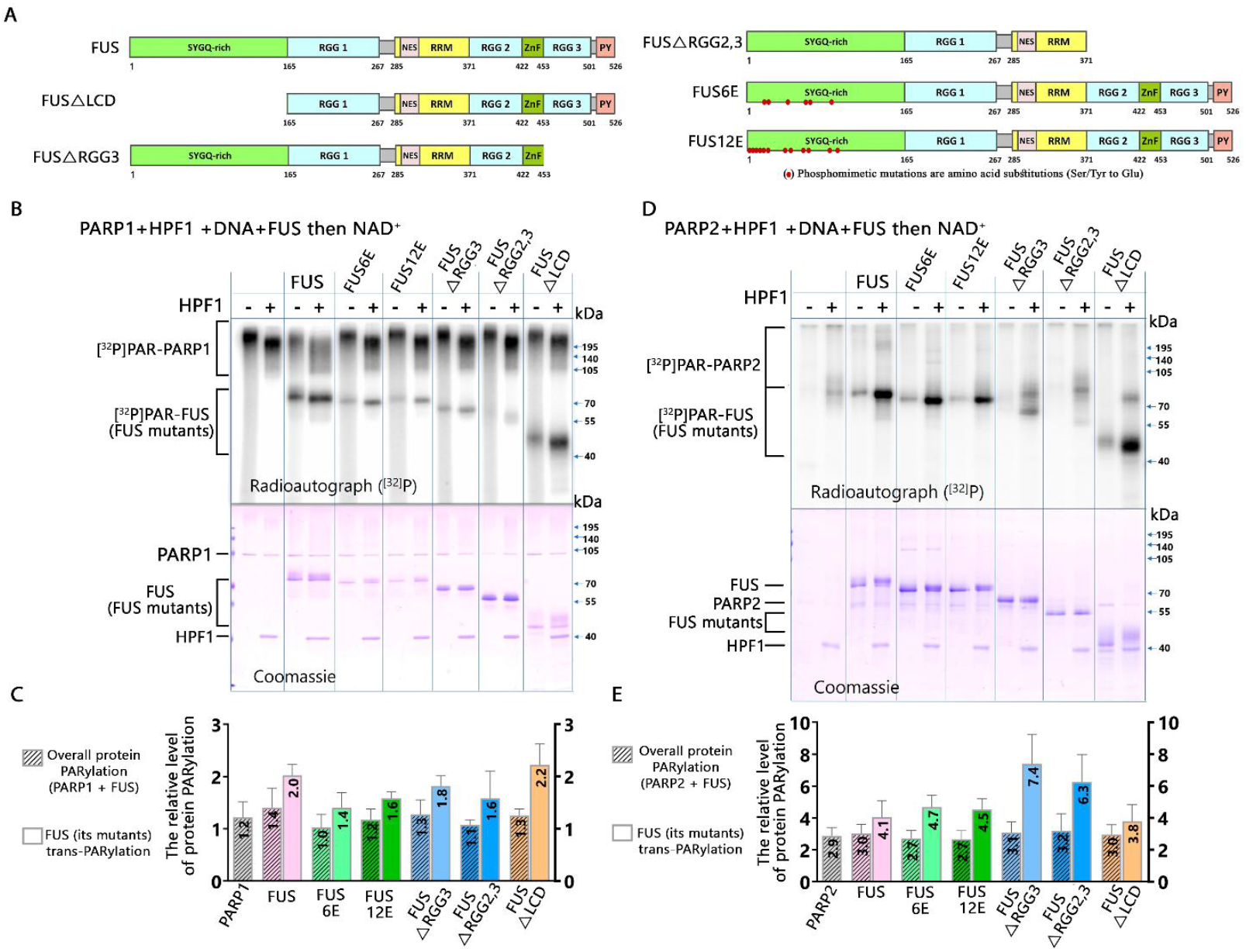
HPF1 affects compartment formation by stimulating PARylation of FUS. (**A**) Domain structure of FUS wild type and its mutants: FUSΔLCD, FUSΔRGG3, FUSΔRGG2,3, FUS6E and FUS12E. (**B**) PARylation of PARP1 (250 nM), FUS and its mutants (1 μM) in the presence of damaged DNA (12.5 nM), HPF1 (1 µM) and 0.3 mM NAD^+^, 0.4 μCi [^32^P]NAD^+^ detected by 10% SDS-PAGE with subsequent phosphorimaging and Coomassie staining. (**C**) The diagrams show the relative levels of protein PARylation (PARP1, FUS or FUS mutants) (the mean ± SD of three independent experiments) from (**B**). Bars filled with dashes represent total protein PARylation in each sample, and bars without dashes represent hetero -PARylation of FUS and its mutants. The bars with dashes represent the total amount of PARylation in each sample, while the bars without dashes represent the amount of hetero-PARylation of FUS or its mutants (FUS6E, FUS12E, FUSΔRGG3, FUSΔRGG2,3, and FUSΔLCD). Relative protein PARylation levels were normalized to the PARylation data in the absence of HPF1. (**D**) PARylation of PARP2 (250 nM), FUS and its mutants (1 μM) in the presence of damaged DNA (12.5 nM), HPF1 (1 µM) and 0.3 mM NAD+, 0.4 μCi [^32^P]-NAD+ detected by 10% SDS-PAGE with subsequent phosphorimaging and Coomassie staining. (**E**) The diagrams show the relative levels of protein PARylation (PARP2, FUS, and FUS mutants) from three independent experiments. The bars with dashes represent the total PARylation of each sample, while the bars without dashes represent the hetero-PARylation of FUS or its mutants (FUS6E, FUS12E, FUSΔRGG3, FUSΔRGG2,3, and FUSΔLCD). The relative level of protein PARylation was normalized to their PARylation level observed in the absence of HPF1.

It should be noted that in contrast to PARP1, we were unable to image the formation of compartments with AFM when FUS was added before PARP2 activation (data not shown). This was possible due to the low level of PAR synthesis catalyzed by PARP2, coupled with a high level of FUS PARylation under these conditions that has to destroy compartment formation (Figure S4B). It is interesting that, unlike PARP1, HPF1 had a stronger stimulatory effect on the level of PARP2-mediated PARylation of FUS wild-type and its mutants (Figure 5D, F). The level of FUS PARylation and its mutants catalyzed by PARP2 were found to be significantly higher (four-to six-fold) in the presence of HPF1 than in its absence (Figure 5E). The stimulation of FUS PARylation by PARP2 in the presence of HPF1 should destroy the compartment formation.

Thus, PARP1 and PARP2 exhibit similar tendencies towards the PARylation of FUS mutants compared to the wild-type FUS: lower levels of modification for FUS6E and FUS12E; higher levels for FUSΔLCD; and very low levels for FUSΔRGG2,3 (Figure 5). These data indicate that HPF1 strongly stimulates FUS hetero-PARylation. Furthermore, deleting the two PAR-binding domains (RGG2 and RGG3) in FUS drastically impairs PARP1- and PARP2-mediated FUS modifications in both the presence and absence of HPF1. As mentioned above, the N-terminal LCD mainly promotes FUS’s self-assembly, interaction with other proteins, and phase separation (44, 62, 63, 64), whereas RGG2,3 domains are involved in the interaction with PAR (9, 30, 36). Thus, the deletion of the RGG(2,3) domains may impair the binding of FUS to PAR covalently attached to PARP1 or PARP2, which may influence PARylation of FUS. The LCD mutations decrease the PARylation of FUS phosphomimetics (FUS6E or FUS12E), which suggests that controlling FUS phase separation could depend on both PARylation and phosphorylation.

Since a shortening of the PAR length was observed in the presence of HPF1 (Figure S3), its stimulatory effect on the level of FUS and its mutants PARylation by PARP1 and PARP2 may be related to an increase in the number of modification sites on this protein. In order to test this hypothesis, PARG treatment was carried out after PARylation of the proteins in the presence of HPF1 (Figure S5). Evaluating the [^32^P]-FUS-ADP-ribose signal after PARG treatment revealed that FUS and its mutants were modified by PARP1 and PARP2 more effectively in the presence of HPF1 than in its absence (Figure S5).

These data suggest that HPF1 stimulates the overall level of FUS PARylation as a result of an increase in modification sites. To determine whether FUS and its mutants were predominantly PARylated on serine residues in the presence of HPF1, the modified proteins were treated with hydroxylamine, which disrupts the ester bond between the terminal ADP-ribose unit and carboxyl group of glutamate and aspartate residues (Figure S5) (65, 66). After treatment with hydroxylamine, the higher level of PARylation of proteins by PARP1 was maintained in the presence of HPF1 and is even stronger for PARP2, indicating a preferential PARylation of FUS and its mutants at serine residues in the presence of HPF1 (Figure S5C, D).

Thus, adding FUS before the addition of NAD^+^ for PARP1(2) activation, i.e., prior to PARP1(2) PARylation, may inhibit compartment formation due to increased FUS PARylation and the covalent attachment of PAR to amino acid residues in RGGdomains. HPF1 may, therefore, play an important role in the regulation of FUS PARylation by PARP1 and PARP2 and thus can contribute to the regulation of compartment formation. The precise regulation of PAR levels and the selection of amino acid residues as ADP-ribose acceptors in proteins may be crucial for coordinating liquid-liquid phase separation at sites of DNA damage and forming DNA-containing compartments with the correct composition and structure.

## DISCUSSION

Chromatin structure is extensively reorganized in response to DNA damage, and the synthesis of PAR catalyzed by PARP1 is one of the fastest damage signals, which serves to coordinate multiple DNA-dependent processes such as DNA repair both during and after stress (1). The rapid recruitment and autoPARylation of PARP1 or PARP2 in response to DNA damage raises the possibility that it serves as a nucleation site for the formation of a DNA repair condensate (3, 12, 22, 30, 31). It is thought that FUS, which regulates the metabolism of mRNA, plays an important role in the formation of biomolecular condensates in a living organism (4, 5, 37, 38, 39, 40, 41). In particular, the ability of FUS to condense and form phase-separated assemblies is influenced by its binding to PAR during PARP1 activation on damaged DNA (9, 30, 49, 50). To date, a wealth of data has revealed functional interactions between FUS, PARP1, and PAR synthesis. For instance, the formation of FUS-rich assemblies in regions of DNA damage after laser microirradiation is dependent on PARP1 activation (29, 30, 31). FUS has been shown to interact with PAR and PARP1, thereby regulating PARP1-dependent PAR synthesis in HeLa cells following genotoxic stress (49). The interaction between FUS and PARylated PARP1 appears to play a specific role in the formation of DNA-rich compartments in vitro (9). Furthermore, PAR synthesis contributes to the recruitment of FUS to sites of DNA damage and promotes its liquid-liquid phase separation in cells (30, 31). Although numerous studies support the importance of PARP1 activity for protein condensation in the nucleus (4, 5), the role of PARP2 in this process has yet to be studied. Both PARP1 and PARP2 are primary sensors of DNA breaks and are involved in base excision or single-strand break repair (10, 11, 52). PARP1 and PARP2 primarily act as sensors of DNA strand breaks, forming DNA repair foci via local synthesis of PAR at sites of DNA damage, loading proteins involved in DNA base excision repair and single-strand break repair, such as scaffold protein XRCC1, PNKP (DNA polynucleotide kinase phosphatase), and DNA polymerase β (3, 12). HPF1 could also be involved in the regulation of ADP-ribosylation-dependent formation of DNA repair foci, such as histone ADP-ribosylation by PARP1 followed by chromatin unfolding and recruitment of repair proteins (22, 23).

The structure of PARP1 and PARP2 is significantly different due to the absence of zinc fingers in PARP2 (1, 10). PARP1’s zinc fingers play an important role in the compartmentalization of PARP1 with DNA in vitro (8, 67), which raises a number of questions about the possibility of the formation of DNA-rich compartments with the participation of PARP2 alone. Indeed, unlike PARP1, PARP2 (due to the absence of zinc fingers) cannot form compartments by dimerization with DNA duplexes, although it can be recruited to compartments by binding to PAR after its synthesis by PARP1 or form Mg^2+^-dependent liquid-like assemblies during auto-PARylation (6, 67). Previously, our analysis of PAR and FUS condensation has been restricted to the analysis of PARP1-FUS interactions (9, 49, 50). The ability of PARylated PARP1 to induce the FUS condensation gave rise to the assumption that PARP2 may also be effective in the regulation of FUS phase separation. Therefore, we investigated the formation of DNA-rich compartments involving FUS and PARylated PARP1 or PARP2 when their activity was regulated with HPF1. The discovery of HPF1, a factor that can regulate PAR synthesis efficiency, PAR length, and the auto- and hetero-ADP-ribosylation activities of PARP1 and PARP2, suggests that HPF1 can potentially influence PAR-dependent FUS phase separation in chromatin. As with PARP1, we found that FUS and PARylated PARP2 form DNA-rich compartments(Figure 2). Thus, PAR synthesis catalyzed by PARP2 alone is also sufficient to form compartments of such type (Figure 2A, C). We also demonstrate that HPF1 can modulate the synthesis of PAR catalyzed by PARP1 and PARP2 in the presence of plasmid DNA. HPF1 reduces the amount of PAR, the size of PARylated molecules, and the size of DNA-rich compartments when FUS was added after PARP1 activation, as it was clearly observed in the case of PARP1 (Figures 1B, C). In the case of PARP2, HPF1 diminishes the level of its auto-PARylation in a certain range of their molar ratio (Figure 2D); however, HPF1 does not significantly affect the size of either PARylated molecules or FUS compartments. (Figure 2B, C). Thus, HPF1 had a lower effect on the formation of DNA-rich compartments with participation of FUS and PARylated PARP2 when FUS was added after the PARylation reaction started (Figure 2C). Thus, despite the presence of HPF1 and the shortening of PAR synthesized by PARP1(2), smaller compartments were formed (Figures 1 and 2). We also demonstrate that PARG efficiently hydrolyzes PAR when PARP1- or PARP2-catalyzed PAR synthesis occurred in the presence of HPF1 and after adding FUS (Figure 3). This appears to correlate with the disruption of FUS compartments observed in the presence of PARG (Figure 3 A, B). Given the importance of PARG-dependent PAR degradation in the translocation of FUS to cytoplasm under oxidative stress (9, 68)-+, one might expect that PARG to be directly implicated in the regulation of the disruption of condensate formation with the participation of FUS and PARylated PARP1 or PARP2. To explore the possible relationship between FUS PARylation and condensation, we investigated compartment formation when FUS was added prior to PARP1(2) activation (Figures 4 and 5). Indeed, FUS was effectively PARylated when it was added before NAD^+^, i.e., prior to the initiation of PARP1(2) PARylation activity (Figures S4 and 5). The current data show that FUS undergoes ADP-ribosylation modification at serine residues under genotoxic stress, suggesting that HPF1 can be involved in regulating FUS PARylation (60, 61). Stimulation of hetero-PARylation of FUS and its mutants was also shown in the presence of HPF1 (Figure 5B, D), and it is worth noting that the stimulation effect was much stronger for PARP2 compared to PARP1 (Figure 5C, E). Using AFM imaging, we were unable to visualize the formation of FUS compartments with PARylated PARP2 under conditions when FUS was added before the addition of NAD^+^ for PARP2 activation, in either the presence or absence of HPF1. It is possible that FUS hetero-PARylation was relatively high under such conditions and FUS interaction with PAR was destroyed, particularly when HPF1 was present (Figures S4 and 5D, E). Thus, HPF1 can switch PARylation from the autopoly(ADP-ribosyl)ation of PARP1(2) to the hetero-PARylation of FUS (Figure 5B, D). These data are in agreement with intensive hetero-PARylation of histones, which was observed for PARP2 but not for PARP1 in the presence of HPF1 (21).

Recent data indicate that HPF1 binds to PARP1(2) at the stage of initiation of PARylation of PARP’s at serine residues and thereby significantly increases the rate of addition of the first ADP-ribose to Ser compared to the rate observed for addition of ADP-ribose to Glu(Asp) in the absence of HPF1 (18). In the presence of HPF1, the elongation reaction of poly(ADP-ribose) synthesis was shown to be inhibited by steric block at the acceptor site (18), and this may explain why we observe a higher number of ADP-ribose residues added to FUS in the presence of HPF1 observed after PARG treatment (Figure S5).

Thus, since HPF1 significantly increases the rate of initiation of PARylation at serine residues, this may lead to enhanced hetero-PARylation of target non-histone proteins such as FUS and thus possibly to regulate compartment assembly and DNA repair (Figure 6). Considering that serine residues in FUS are primarily found in the N-terminal LCD domain, which are crucial for the FUS phase transition (31, 44, 62), the formation of DNA-rich condensates may be influenced by PARylation of serine residues in the LCD by PARP1 or PARP2 in the presence of HPF1.

**Figure 6.**
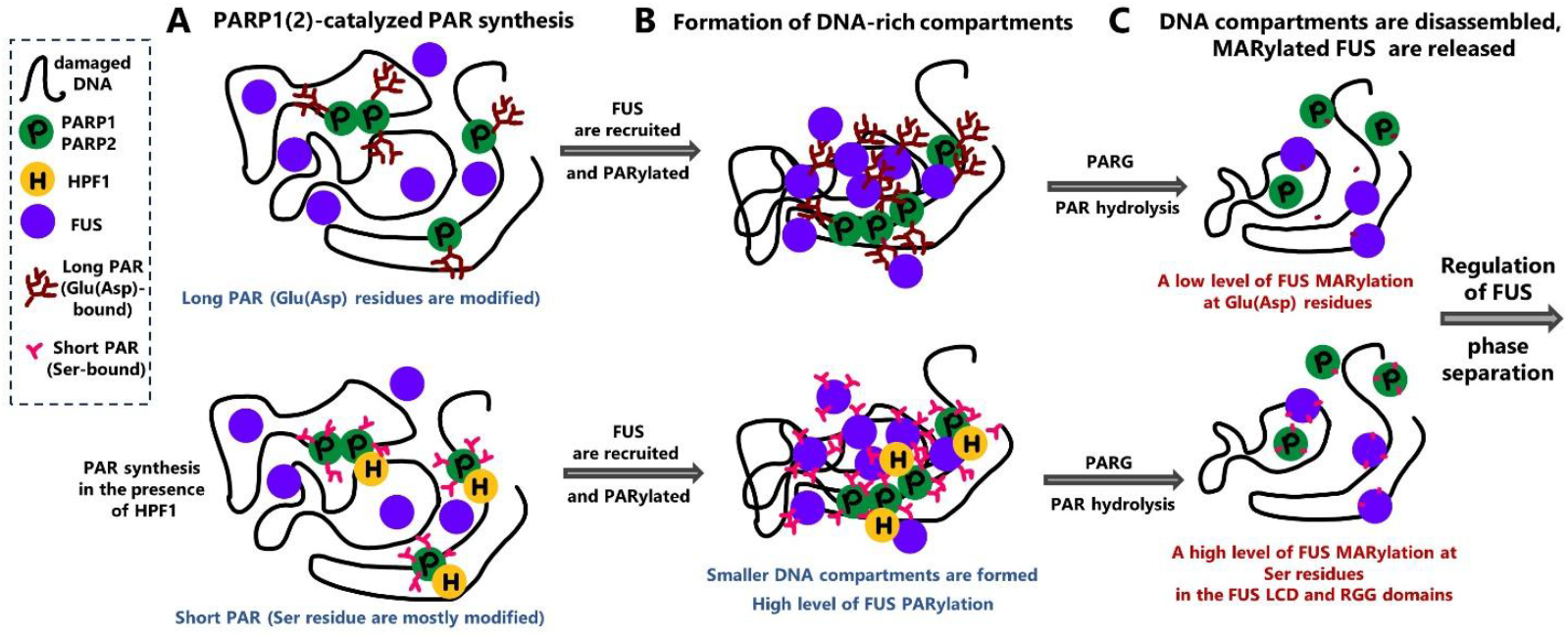
A simplified model of HPF1-dependent regulation of formation of DNA-rich compartments when protein PARylation occurs in the absence of HPF1 or presence of HPF1. (**A**) PARP1- or PARP2-dependent synthesis of long or short PAR in the absence or presence of HPF1, respectively. (**B**) Formation of DNA-rich compartments. In the absence of HPF1, a moderate level of FUS PARylation proceeds. FUS PARylation increases significantly when HPF1 is present. (**C**) Disassembly of DNA-rich compartments due to PARG-dependent PAR degradation. FUS has MARylated glutamate (aspartate) residues when HPF1 is absent. In the presence of HPF1, FUS is more intensively MARylated at serine residues. The serine-ADP-ribosylation could influence FUS phase separation and its participation in the formation of biomolecular condensates.

Due to the significant increase in the level of FUS PARylation in the presence of HPF1, we can expect a stronger stress response. FUS is an important transcription factor, and PARylation is necessary for its translocation into the cytoplasm, entry into stress granules, and regulation of translation under stress conditions (9, 68). It is worth noting that it has been shown that the regulation of stress granule formation in certain diseases such as amyotrophic lateral sclerosis depends on the activity of PARP1 and requires further study (68). Recent studies show that HPF1 impacts PARP1 retention and increases the rate of PARylation initiation events of serine residues at the initial time point, but with inhibition of elongation (decrease in PAR length) (18, 19). PARP2 is known to be less active compared to PARP1 (10, 11); however, HPF1 similarly stimulates PARP2 activity at the initiation stage, which ultimately may result in strong stimulation of PARP2 observed in our experiments (Figure 4F).

Recent data suggest that condensate formation may coordinate the following processes: opening of chromatin, recruitment or exclusion of specific proteins, concentration and fixation of several DNA lesions at one site, and enhancement or delay of regulatory signals until condensate disassembly after DNA repair (40, 43). In this context, the study of the HPF1 effect as a regulator of PAR synthesis, serine PARylation of acceptor proteins by PARP1 and PARP2, and the formation of FUS compartments should be fully explored. This is important to understand the overlapping of PARP1 and PARP2 functions evidenced by the capacity of cells and animals to survive after knocking out one of these two important enzymes (52, 69), but also for the perspective of developing PARP1 or PARP2 selective inhibitors to lower the toxicity of unspecific inhibitors (70).

## DATA AVAILABILITY

The original contributions presented in this study are included in the article and Supplementary Materials; further inquiries can be directed to the corresponding author.

## SUPPLEMENTARY DATA STATEMENT

Supplementary Data are available at NAR Online.

## ACKNOWLEDGEMENTS

The authors are grateful to T. A. Kurgina (ICBFM SB RAS, Novosibirsk), N. S. Dyrkheeva (ICBFM SB RAS, Novosibirsk), and S. I. Shram (Institute of Molecular Genetics RAS, Moscow) for PARG, APE1 and NMNAT, respectively. The authors gratefully acknowledge UEVE Université Paris-Saclay and Genopole EVRY for the constant support of the Laboratoire Structure-Activité des Biomolécules Normales et Pathologiques (INSERM U1204, University of Evry/Paris-Saclay), as well as the INSERM PRI grant [RaPiD].

## AUTHOR CONTRIBUTIONS STATEMENT

Conceptualization, O.I.L. and D.P.; methodology, M.V.S., A.S.S., and L.H.; validation, A.S.S. and M.V.S.; formal analysis, A.S.S. and M.V.S.; investigation, A.S.S., K.M.M., and L.H.; resources, O.I.L. and D.P.; writing—original draft preparation, M.V.S. and A.S.S.; writing—review and editing, O.I.L., D.P., and M.V.S.; visualization, A.S.S. and M.V.S.; supervision, O.I.L. and D.P.; project administration, O.I.L. and D.P.; funding acquisition, O.I.L. and D.P. All authors have read and agreed to the published version of the manuscript.

