## Supplementary Figures for "HPF1 REGULATES THE FORMATION OF FUS-DEPENDENT COMPARTMENTS BY PARP1 AND PARP2 ACTIVATION ON DAMAGED DNA": SUPPLEMENTARY DATA.pdf

#### SUPPLEMENTARY FIGURES

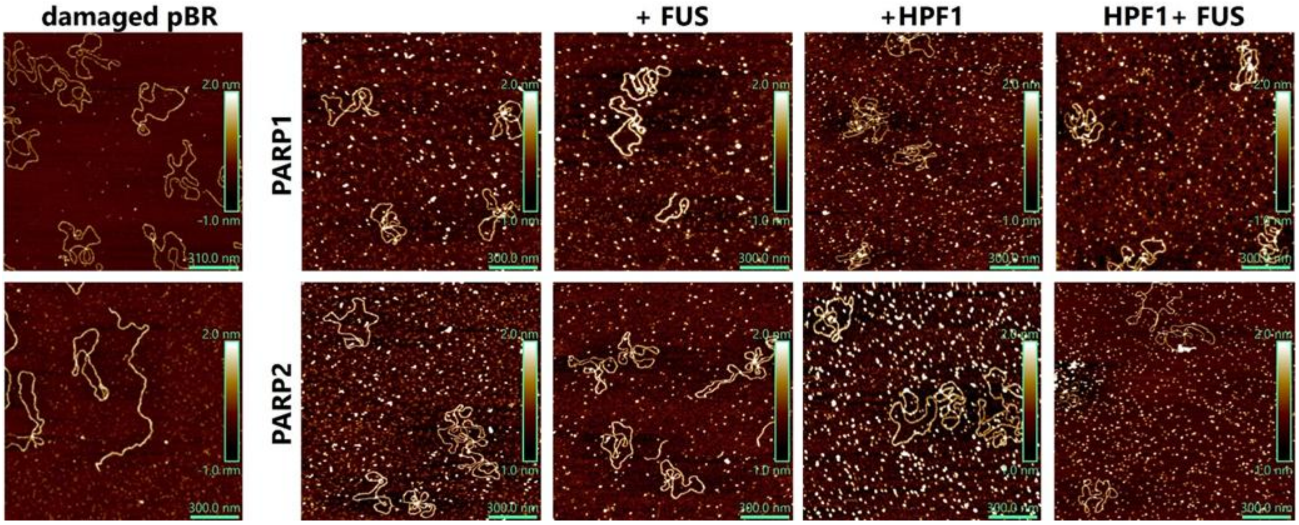


**Figure S1.** AFM images of damaged pBR in the absence or presence of PARP1, PARP2, HPF1, FUS before adding NAD$^+$ and initiation of PAR synthesis. pBR plasmid (12.5 nM) was incubated with PARP1(2) (30 nM), HPF1 (120nM) or FUS (400 nM) as indicated at the figure legends, deposited on mica surface imaged by AFM in air.

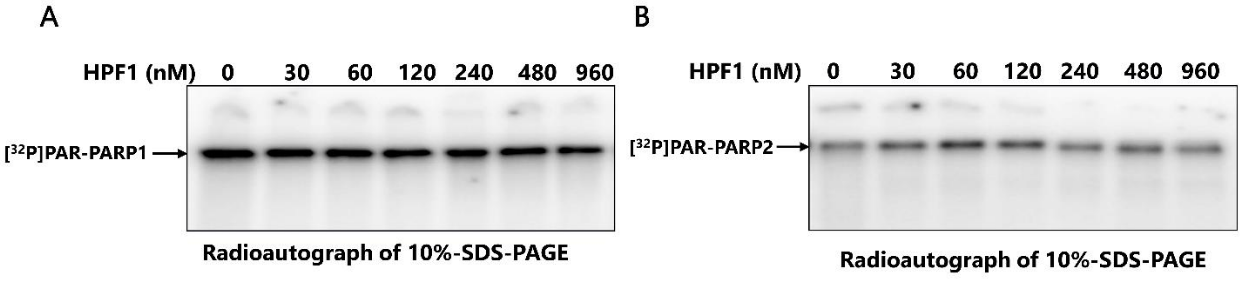


**Figure S2.** PARP1 (A) or PARP2(B) auto-PARylation detected after SDS-PAGE with phosphorimaging. An in vitro poly(ADP-ribosyl)ation assay was performed in the reaction mixtures (20 µL) contained 12.5 mM HEPES pH 7.6, 12.5 mM NaCl, 1 mM DTT, 5 mM $MgCl_2$, 12.5 nM damaged DNA, 30 PARP1(2), 30-960 nM HPF1, 0.3 mM NAD$^+$, 0.4 µCi [$^{32}$P]-NAD$^+$. The reactions were initiated by the addition of NAD$^+$. The reaction mixtures were incubated at 37 °C for 15 min for PARP1 and 30 min for PARP2 at 37$^0$C and stopped by adding SDS-sample buffer and heating for 5 min at 90°C. The reaction mixtures were analyzed by 10% SDS-PAGE with subsequent phosphorimaging.

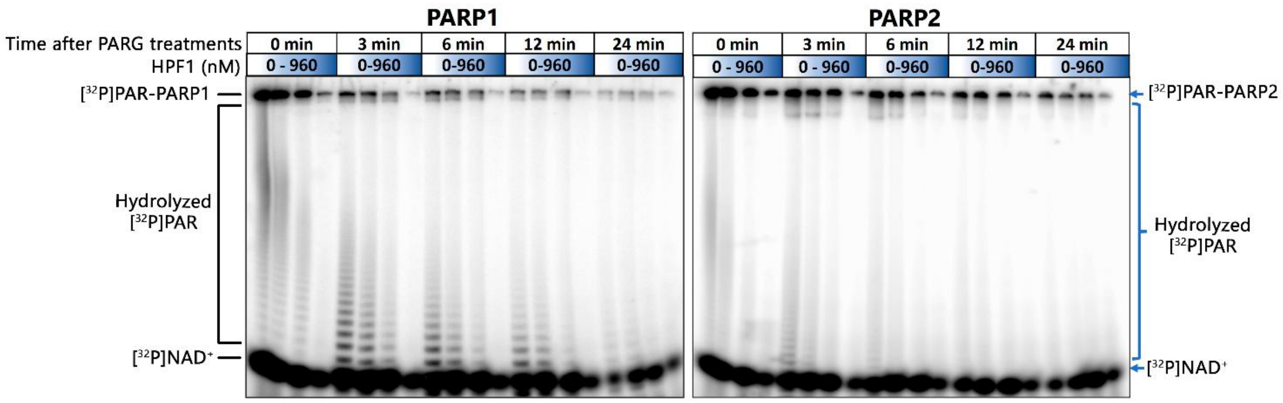


**Figure S3.** PAR degradation by PARG after the addition of FUS.

For the analysis of the PARylation of PARP1(2) after PARG treatment, 30 nM PARP1(2), 400 nM FUS, 12.5 nM damaged pBR plasmid and 30-960 nM HPF1 (as indicated on figure legend) were incubated with 0.3 mM $NAD^+$ and 0.4 µCi [$^{32}$P]-$NAD^+$ at 37 °C for 15 min for PARP1 and 30 min for PARP2 at 37$^0$C and then were stopped by Olaparib to a final concentration of 250 nM. Then, PARG was added to a final concentration of 40 nM, after which the mixture was incubated for 3-24 minutes at 37°C. The reaction mixture (20 µL) was stopped by adding 4 µL of the loading solution containing 90% formamide, 50 mM EDTA, 0.1% xylene cyanol, and 0.1% bromophenol blue, heated for 5 min at 95°C, and the products were separated by denaturing electrophoresis in 10% polyacrylamide gel followed by visualization with phosphorimaging.

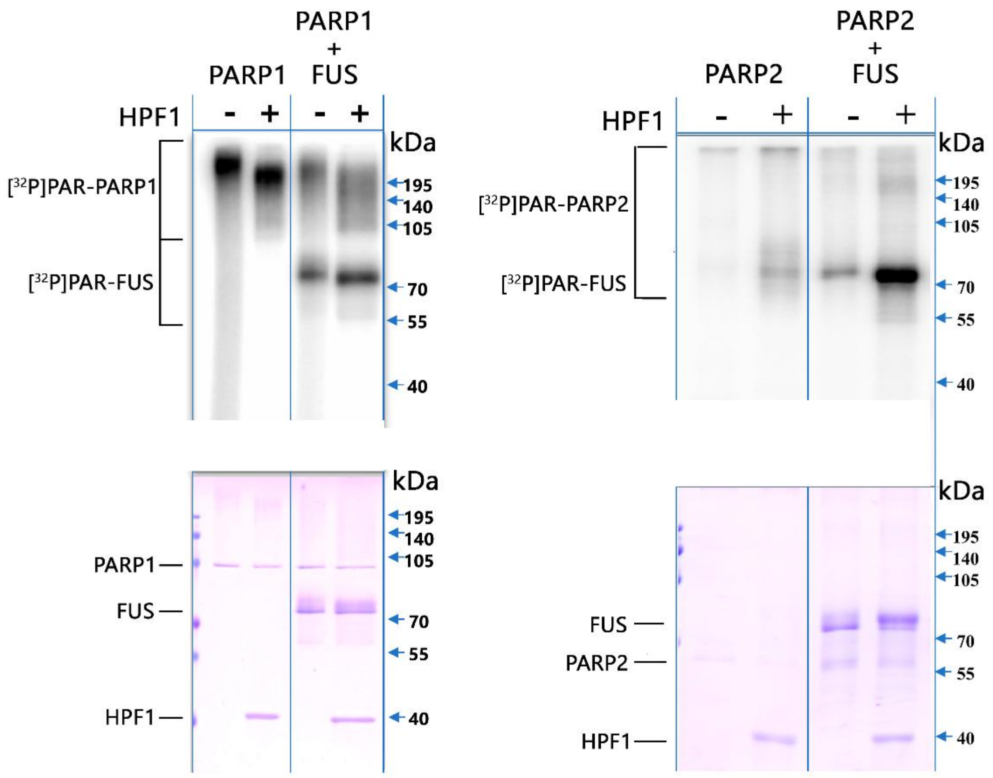


**Figure S4.** HPF1 stimulates hetero-PARylation of FUS by PARP1 and PARP2. PARylation of PARP1 or PARP2 (250 nM), FUS (1 µM) in the presence of damaged DNA (12.5 nM), HPF1 (1 µM) and 0.3 mM $NAD^+$, 0.4 µCi $[^{32}P]NAD^+$ detected by 10% SDS-PAGE with subsequent phosphorimaging and Coomassie staining.

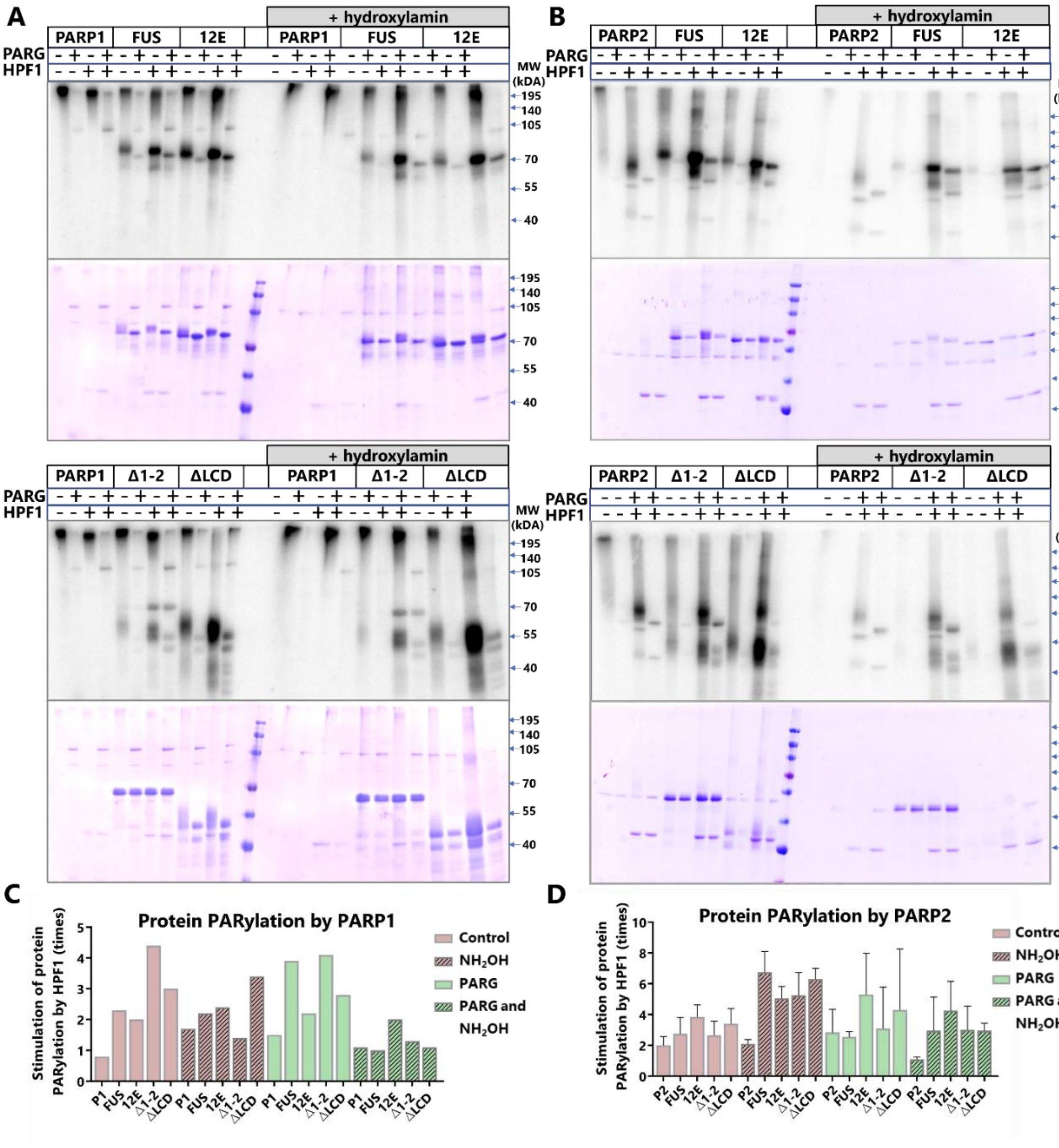


**Figure S5.** Treatment with hydroxylamine and PARG shows that FUS and its mutants are preferentially PARylated at serine residues, and that the number of protein modification sites increases in the presence of HPF1.

**(A)** PARylation of PARP1 (250 nM), FUS or its mutants (1 µM) in the presence of damaged DNA (12.5 nM), HPF1 (1 µM) and [$^{32}$P]-NAD$^+$ (300 µM) followed by PARG and/or hydroxylamine treatment. PARylation proteins were detected by 10% SDS-PAGE with subsequent phosphorimaging and Coomassie staining.

**(B)** PARylation of PARP2 (250 nM), FUS or its mutants (1 µM) in the presence of damaged DNA (12.5 nM), HPF1 (1 µM) and [$^{32}$P]-NAD$^+$ (300 µM) followed by PARG and (or) hydroxylamine treatment. PARylation proteins were detected by 10% SDS-PAGE with subsequent phosphorimaging and Coomassie staining.

**(C)** The diagram shows the relative levels of PARylation of PARP1 and hetero-PARylation of FUS (or its mutants) before and after treatment with PARG or (and) hydroxylamine. The relative levels of protein PARylation were normalized to the protein modifications observed in the absence of HPF1 from (A) (the mean ± SD of two independent experiments).

**(D)** The diagram shows the relative levels of PARylation of PARP2 and hetero-PARylation of FUS (or its mutants) before and after treatment with PARG or (and) hydroxylamine. The relative levels of protein PARylation were normalized to the protein modifications observed in the absence of HPF1 from (B) (the mean ± SD of two independent experiments).

### SUPPLEMENTARY METHODS

#### PARG and Hydroxylamine treatment

Protein poly(ADP-ribosyl)ation assay was performed in the reaction mixtures (20 µL) contained 12.5 mM HEPES pH 7.6, 12.5 mM NaCl, 1 mM DTT, 100 mM urea, 5 mM $MgCl_2$, 0.3 mM $NAD^+$, 0.4 µCi [$^{32}P$]-$NAD^+$, 12.5 nM damaged DNA, 250 nM PARP1(2), 1 µM HPF1, 1 µM FUS or its mutants as indicated at the figure legends. The reactions were initiated by the addition of $NAD^+$ and incubated at 37°C for 15 min for PARP1 and 30 min for PARP2 at $37^0$C, after that stopped by the addition of olaparib to a final concentration of 250 nM. PARG was added to a final concentration of 40 nM and then the reactions were incubated for 24 min at 37°C. Where indicated, the reactions were treated with hydroxylamine ($NH_2OH$, pH 7.5) to a final concentration of 1 M and then incubated for 1 h at 37 °C.

The reactions were stopped by the addition of SDS sample buffer and by heating for 2 min at 95 °C. The products were separated by denaturing 10% PAGE (with SDS) with subsequent phosphorimaging and colloidal Coomassie staining.
